# Comparative Effects of Hypoxic vs. Normoxic Mesenchymal Stem Cell-Derived Extracellular Vesicles on Tissue Repair Following Volumetric Muscle Loss (VML)

**DOI:** 10.64898/2025.12.31.697216

**Authors:** Avantika Jain, Amelia Ridolfo, Mathangi Mohankumar Subramaniam, David Johnson, Jacki Kornbluth, Koyal Garg

## Abstract

Volumetric muscle loss (VML) is an irreversible muscle injury that results in chronic functional impairment. Mesenchymal stem cell (MSC)-derived extracellular vesicles (EVs) can facilitate tissue repair through immunomodulatory, angiogenic, and anti-fibrotic effects. However, their low yield and poor on-site retention limit their therapeutic efficacy. Hypoxia can boost MSC metabolism, proliferation, and EV production. Hypoxic (3% O_2_) preconditioning of MSCs increased the yield of EVs (30-300 nm) by 1.5-fold but decreased the expression of characteristic EV markers (i.e., CD81, ICAM, and FLOT1). Fibrin hydrogels promote skeletal muscle regeneration and can sequester EVs via integrins or electrostatic interactions. We hypothesized that encapsulating EVs in fibrin hydrogels would further enhance regeneration and prolong the retention of EVs at the VML injury site. VML was created by removing ∼20% of the gastrocnemius-soleus muscle’s mass in mice using a 3 mm biopsy punch. EVs (4.48×10^10^ particles/mL) derived from MSCs cultured under hypoxic (Hypo-EV) or normoxic (Norm-EV) conditions were encapsulated within fibrin hydrogels and implanted at the VML injury site. Fibrin hydrogels containing PBS (PFG) were used as controls. On day 14 post-injury, Norm-EV treatment resulted in increased muscle mass, angiogenesis, and myofiber regeneration relative to the Hypo-EV group. Both the Norm-EV and Hypo-EV treatment groups reduced macrophage infiltration at the injury site compared to the PFG. These findings highlight that while both Norm-EV and Hypo-EV exhibit immunomodulatory effects, they differ in their regenerative potential. We speculate that hypoxic conditions could have caused MSCs to prioritize survival over repair-promoting activities, thereby producing EVs with less pro-regenerative signals. The increased quantity of EVs in response to hypoxia doesn’t compensate for their diminished regenerative potential, highlighting the importance of quality over quantity when considering EVs for therapeutic applications.

**Graphical Abstract:** Jain et al., Comparative Effects of Hypoxic vs. Normoxic Mesenchymal Stem Cell-Derived Extracellular Vesicles on Tissue Repair Following Volumetric Muscle Loss (VML)

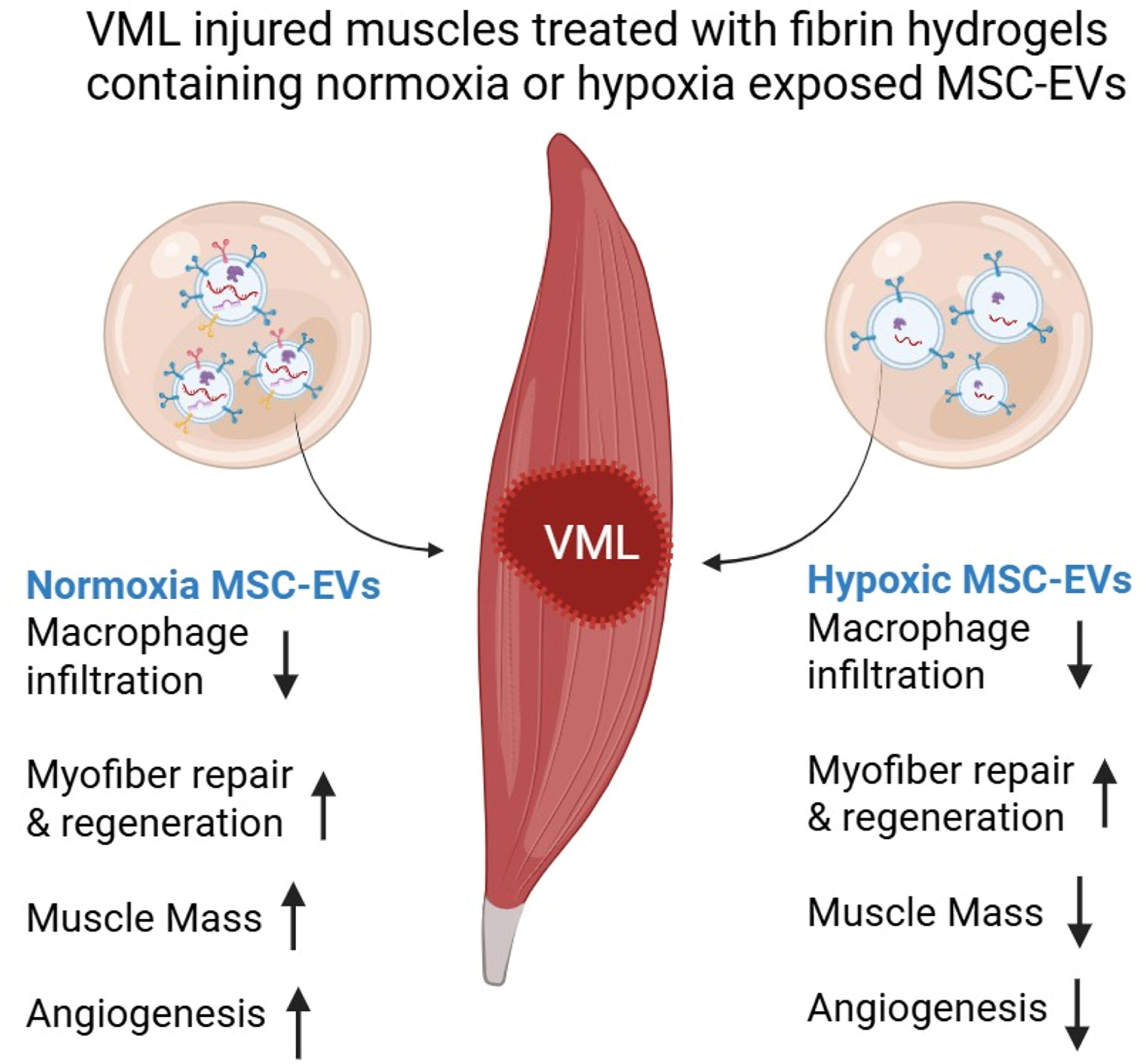

## 1. Introduction

Musculoskeletal conditions represent a major public health burden in the United States, affecting ∼127.4 million individuals and surpassing all other health conditions as the leading driver of national healthcare expenditure at over $381 billion annually [1]. Severe musculoskeletal injuries are frequently complicated by volumetric muscle loss (VML) [2], defined as traumatic or surgical loss of muscle tissue (>20%) with resultant functional impairment [3]. VML injuries result in persistent inflammation, extensive fibrosis, and with little to no muscle regeneration [4–9]. VML has emerged as a critical limiting factor in limb salvage [10]. The inability to restore functional muscle mass often renders a salvaged limb non-functional, which may ultimately require delayed amputation. Within the military healthcare system, over 90% of medical retirements involving muscle-related disability are linked to VML [11]. Current clinical treatments, such as physical therapy and bracing, are focused mainly on supporting and strengthening the remaining muscle mass post-VML and not on regenerating the lost muscle tissue [12, 13]. Muscle reconstruction methods, such as autologous muscle flap surgery [14] or transplantation of composite musculoskeletal tissue graft [15] pose challenges such as donor site morbidity, tissue rejection, lifelong immunosuppression, and possible infections.

Skeletal muscle tissue engineering therapies, which typically involve the use of stem cell populations with myogenic potential and biologic scaffolds composed of extracellular proteins, have emerged as a promising approach to treat VML [15, 16]. Bone marrow-derived mesenchymal stem cells (BM-MSCs) are a multipotent, adult stem cell population that can reliably differentiate into bone, cartilage, and adipose tissues [17, 18]. Several studies have used BM-MSCs with various biomaterials to treat VML. In small animal models of VML, MSCs have been delivered through direct intramuscular injection [19], fibrin [20] or fibrin-laminin hydrogels [21], extracellular matrix (ECM) scaffolds [22] or platelet-rich plasma-derived fibrin microbeads [23]. In these studies, BM-MSCs promoted muscle recovery post-VML through various mechanisms, including immunomodulation, angiogenesis, anti-fibrotic action, and myofiber regeneration. However, the BM-MSCs rarely differentiated directly into myofibers. BM-MSCs exert these pro-regenerative effects primarily through paracrine activities, such as the secretion of growth factors, cytokines, chemokines, and extracellular vesicles (EVs) [24, 25]. Therefore, the therapeutic benefits of MSCs may be realized by focusing on their secretome. Of particular interest, EVs are biological nanovesicles secreted by mammalian cells for intercellular communication [26]. These vesicles contain bioactive molecules, such as lipids, proteins, mRNAs, and microRNAs (miRNAs), which are unique to the cell of origin [27, 28]. As an acellular MSC byproduct, EVs can readily circulate through organs, elicit a minimal immune response, avoid phagocytosis, and elicit cellular responses by binding to specific receptors on the target cell [29]. As a therapeutic approach, EV delivery circumvents any safety concerns associated with stem cell expansion, differentiation into desired lineages, and tumorigenesis [30]. Moreover, EVs are stable for 6 months if stored at -20°C in an isotonic buffer [31] and represent a promising “off-the-shelf” treatment for inflammatory or fibrotic conditions [32]. In acute [33, 34] and chronic [35] models of muscle injury, MSC-EV delivery has resulted in improved outcomes.

However, the practical application of EVs for human population-based clinical treatments is severely limited due to insufficient cellular production of EVs [36]. Previous studies have shown that hypoxic preconditioning (HP) makes cells resistant to ischemic damage. It enhances stem cell transplantation through autocrine and paracrine signaling, increases survival rate, and angiogenesis [37–41]. Studies have also shown that HP not only triggers an increase in the release of EV [18, 42], but also enhances their potential for tissue repair [34, 43–49].

To improve the retention of EVs at the VML injury site, MSC-EVs derived from hypoxic and normoxic cultures were encapsulated within fibrin hydrogels and implanted into the defect. Fibrin hydrogels have been used extensively for skeletal muscle tissue engineering applications by us and others [21, 50–52]. Additionally, fibrin hydrogels have been successfully used to deliver EVs into injured tissues in previous studies [53–56]. Therefore, we hypothesized that (1) HP of MSCs will increase EV production, and (2) MSC-EVs isolated from hypoxic and normoxic cultures will have distinct effects on immunomodulation and muscle regeneration.

## 2. Materials and Methods

### 2.1. Cyclic pre-conditioning of BM-MSCs in Hypoxia or Normoxia

Murine bone-marrow derived mesenchymal stem cells (BM-MSCs) (from Cyagen Biomodels LLC., Santa Clara, CA, USA), passage 4-6, were cultured in MSC culture media containing Dulbecco’s Modified Eagle Medium: Nutrient Mixture F-12 (DMEM/F-12) (Cytiva Life Sciences, Marlborough, MA, USA, Catalog No. SH30023.FS) supplemented with 10% exosome-free fetal bovine serum (FBS) (Thermo Fisher Scientific, Waltham, MA, USA, Catalog No. A2720803) and 1% penicillin/streptomycin (P/S) (Corning Inc., New York, N.Y., USA, Catalog 30-002-CI). The BM-MSCs used in this study have been previously characterized in our lab. They express CD146 and SCA-1 in culture and can differentiate into adipocytes, chondrocytes, and osteocytes [21]. BM-MSCs were seeded at a cell density of 5 million cells/ flask in two identical T-75 cm^2^ tissue culture flasks (Thermo Fisher Scientific, Waltham, MA, USA, Catalog No. FB012937), flask-1 and flask-2. BM-MSCs were allowed to attach to the flask surface for 24 hours under optimal cell culture conditions (37°C, 5% CO_2_, 21% O_2_). The culture medium in both flasks was then replaced with fresh 10 ml MSC culture medium. BM-MSCs in flask-1 were cultured in normal oxygen conditions (normoxia) (21% O_2_). BM-MSCs in flask-2 were maintained in low oxygen conditions or hypoxia (3% O2) using a hypoxia chamber (STEMCELL Technologies) flushed with mixed gas composed of 3% O_2_, 5% CO_2_, 92% N_2_. The hypoxia chamber was re-purged at 24 hours according to manufacturer’s instructions to maintain the hypoxia environment for 48 hours. Healthy cell morphology was confirmed through visual inspection. Post this, conditioned medium (CM) was collected from both flasks and cells were replenished with fresh medium to facilitate cell recovery for 24 hours under normoxic conditions. The respective normoxic (for flask-1) and hypoxic (for flask-2) exposures were repeated two more times for 48 hours each, separated by 24 hours recovery period under normoxia. Thus, a total of 3 such pre-conditioning cycles of normoxia or hypoxia were performed, and CM was collected at the end of each cycle and pooled for extracellular vesicle (EV) isolation.

### 2.2. EV Isolation and Characterization

ExoQuick-TC® Ultra for Tissue Culture Media (System Biosciences, Palo Alto, CA, USA, Catalog No. EQULTRA-20TC-1) was used for EV isolation following manufacturer’s instructions [57]. The CM collected post-normoxic or hypoxic pre-conditioning of BM-MSCs was processed separately for EV isolation. The CM was centrifuged at 3000× g for 15 minutes to remove cell debris. CM supernatants were collected and subjected to protein concentration at 4000× g for 15 minutes using Pierce^TM^ protein concentrators, 3K molecular weight cut-off (MWCO) (Thermo Fisher Scientific, Waltham, MA, USA, Catalog No. 88526). This step was added to reduce media diluent and ensure enhanced action Exo-Quick TC Ultra polymer on EV precipitation from the CM. Exo-Quick TC Ultra polymer was mixed with concentrated CM at a ratio of 1:5 and incubated at 4°C for 48 hours. The samples were centrifuged at 3000× g for 10 minutes to pellet the EVs. The supernatant was discarded, and the pelleted EVs were added to pre-washed Exo-Quick-TC Ultra purification columns provided in the kit. The columns were spun at 1000× g for 30 seconds to elute purified EVs. The purified EVs were suspended in a storage buffer provided in the kit and stored at -20°C until needed.

EVs were characterized based on size and expression of characteristic markers. Nanoparticle Tracking Analysis (NTA; NanoSight Pro, Malvern Panalytical Ltd., Worcestershire, UK) was used to determine the size distribution and concentration (particles/ml) of EVs. The protein concentration of EV sample was measured using a NanoDrop 2000 spectrophotometer (Thermo Scientific, Thermo Fisher Scientific, Waltham, MA, USA) at 280 nm. Using the obtained protein concentration, the EV sample was diluted in 1× sterile phosphate-buffered saline (PBS) to 10 μg/ml. The diluted EV sample was passed through a flow cell via syringe and analyzed using NTA as per the manufacturer’s recommended settings.

Expression of characteristic EV markers was assessed using Exo-Check^TM^ Exosome Antibody Array (System Biosciences, Palo Alto, CA, USA, Catalog No. EXORAY200B-4) following manufacturer’s instructions and reagents provided in the kit. The membrane had the following markers for EV characterization: CD63, CD81, epithelial cell adhesion molecule (EpCAM), annexin 5 (ANXA5), tumor susceptibility gene 101 (TSG101), flotillin 1 (FLOT 1), intercellular adhesion molecule (ICAM), and ALG-2-interacting protein X (ALIX). The membrane also had Golgi marker GM130 to detect cellular contamination, two positive controls, and one blank control [58–60]. The normoxic or hypoxic pre-conditioned BM-MSC-derived EVs (50 μg) were processed separately. EVs were dissociated with lysis buffer and the lysed EV components were labelled using a labelling reagent. The excess labelling reagent was removed using purification columns provided in the kit. The purified labelled EV components were mixed with a blocking buffer, loaded onto the Exo-Check^TM^ Exosome Antibody Array and incubated overnight at 4°C. The next day, membranes were washed 2 times using the washing buffer, then incubated with the detection buffer for 30 minutes. Post this, membranes were washed again 3 times. A chemiluminescent developer (BioRad, Hercules, CA, USA, Catalog No. 1705061) mixture was used to detect the expression of EV markers. The array was imaged using a BioRad ChemiDoc Imaging System (BioRad, Hercules, CA, US). The band intensity of markers on the array was quantified using ImageJ.

### 2.3. In vitro macrophage assay

Macrophages (MΦs) (RAW 264.7, ATCC, Manassas, VA, USA) were cultured with Rosewell Park Memorial Institute (RPMI-1640, Sigma-Aldrich, Saint Louis, MO, Catalog No. 72400-047) media with 10% exosome-free FBS (Thermo Fisher Scientific, Waltham, MA, USA, Catalog No. A2720803) and 1% P/S (Corning Inc., New York, N.Y., USA, Catalog 30-002-CI). MΦs were seeded at a cell density of 100,000 cells/well (n=4/group) in a 48-well plate. The seeded MΦs were stimulated and activated using 50 ng/ml of lipopolysaccharide (LPS) for 24 hours. After activation, the media was replaced and MΦs were incubated with one of three treatments: fresh RPMI media (control group), media containing 25 μg/ml of normoxic pre-conditioned BM-MSC derived EVs (Norm-EVs) or media containing 25 μg/ml of hypoxic pre-conditioned BM-MSC derived EVs (Hypo-EVs). After incubation for 72 hours with respective treatments, cell culture supernatants were collected from each well separately and stored at -20°C for future analysis. The release of trophic factors by MΦs in response to respective treatments was quantified using enzyme linked immunosorbent assays (ELISAs). ELISA kits for interleukin-6 (IL-6; Catalog No. 900-K50K), tumor necrosis factor-alpha (TNF-α, Catalog No.900-M54), and vascular endothelial growth factor (VEGF, Catalog No. 900-K99) were purchased from PeproTech ®, parent company Thermofisher Scientific, Cranbury, NJ, USA. Manufacturer’s instructions were followed to perform the ELISAs.

### 2.4. Fibrin Hydrogel Formation

EV encapsulated fibrin hydrogels (600 μl total per gel) were prepared by combining bovine fibrinogen (Catalog No. F8630), thrombin (Catalog No. T7513), EV sample volume containing 4.498×10^10^ particles/ml, calcium chloride (Catalog No. 21115), and protease inhibitor (Catalog No. P8340). All reagents for hydrogel synthesis were purchased from Sigma-Aldrich (parent company Millipore Sigma, Saint Louis, MO, USA). Plain fibrin gels (PFG) were prepared identically, substituting with an equivalent volume of 1× sterile PBS for the EV sample. Fibrinogen was dissolved in 3.5 mL of 0.9% NaCl solution at a concentration of 50 mg/mL, and a protease inhibitor was added at a dilution of 1:1000 to prevent *in vivo* degradation of the hydrogel. For each hydrogel in a 48-well plate, volume of the fibrinogen solution corresponding to a final fibrinogen concentration of 35 mg/ml was pipetted into a well. EVs (4.498×10^10^ particles/ml) or an equivalent volume of 1× PBS was added to the fibrinogen solution in the well to form EV-encapsulated and PFG, respectively. To each well, 50 μl of calcium chloride (20 mM) was added to facilitate the cleaving of fibrinogen by thrombin and fibrin polymerization to form fibrin hydrogel [61]. Finally, 50 μl of 20 U/mL thrombin was added to the well. The well plate was placed in an incubator at 37°C for 20 minutes to facilitate fibrinogen-thrombin crosslinking to form a hydrogel.

### 2.5. EV Release study

Plain fibrin hydrogels (PFG) or normoxic EV encapsulated hydrogels (Norm-EVs) (n=4/group) were prepared as described above. A 6 mm biopsy punch (Vitality Medical, Salt Lake City, UT, Catalog No. 16-1315) was used to punch out hydrogel disks. The hydrogel disks were placed in separate wells of a 48-well plate. 1× sterile PBS (400 µl) was added on top of each hydrogel disk. The PBS was collected and replaced after every 24 hours for a total of 5 days. The presence of EVs released in the PBS was detected using the ExoCheck^TM^ antibody array (System Biosciences, Palo Alto, CA, USA, Catalog No. EXORAY200B-4) following manufacturer’s instructions as previously mentioned in method section 2.2.

### 2.6. Animal Experiments

All animal work was performed in accordance with the Animal Welfare Act, the Animal Welfare Regulations, and following the Guide for the Care and Use of Laboratory Animals. The procedures were approved by the Saint Louis University Institutional Animal Care and Use Committee (AUP Protocol #2612). Adult (14-16 weeks old) female 129S1/SUMJ mice were purchased from Charles River Laboratory and housed in an Association for Assessment and Accreditation of Laboratory Animal Care International-accredited vivarium. Mice were provided with food and water *ad libitum*. The animals were weighed before surgery and anesthetized using 1-2% isoflurane (Covetrus North America, Dublin, OH, USA). The lower hindlimbs were shaved and aseptically prepared using 10% povidone–iodine (Covetrus North America, Dublin, OH, USA) and 70% isopropyl alcohol (Fisher Chemical, Waltham, MA, USA). A lateral incision in the skin was made, and blunt dissection separated the musculature from the surrounding skin and fascia, revealing the gastrocnemius-soleus (GAS) muscle complex. A metal plate was inserted under the GAS complex, and a 3 mm biopsy punch was used to create a full-thickness, ∼20% muscle mass defect in the belly of the GAS complex [51, 62]. The VML defect was made in the GAS complex of both hindlimbs in a mouse. Muscle biopsies were recorded to ensure consistency. A total of 20 mice were randomly divided into four experimental groups as follows: plain fibrin hydrogel (PFG; n=6), hydrogel encapsulated with Norm-EVs (n=6), and hydrogel encapsulated with Hypo-EVs (n=6). Additionally, a no treatment (NT; n=2) VML injured group was used for comparison. Both hindlimbs of a given mouse received either the same treatment or were left untreated. The hydrogels were prepared on the same day as the surgery. For implantation, hydrogels of 6 mm diameter were punched out of a complete hydrogel in a 48-well plate. The punched hydrogel was carefully removed from the well using sterilized forceps and placed both within and atop the VML defect site. The 6 mm punch was ∼25% of the total gel volume. Thus, we are delivering 25% of the EVs that were initially included in the gel. The hydrogel contained 4.5×10^10^ EV particles/mL; therefore, in a 6 mm punch of the hydrogel, we delivered 1.125×10^10^ EV particles per hydrogel. The skin was pulled over the hydrogel and sutured together using simple interrupted Prolene (5-0; Ethicon) sutures. Buprenorphine (1mg/kg; Wedgewood Pharmacy, Swedesboro, NJ) was administered at the nape of the neck subcutaneously to minimize pain post-surgery. Investigators were blinded to group allocation during data collection, and the animals were identified by numbers.

After 14 days of recovery, animals were euthanized with carbon dioxide inhalation, and GAS muscle tissue was dissected carefully along with any visible non-integrated hydrogel fragments. The harvested GAS muscle tissue and hydrogel fragments (if any) were weighed separately. Post-weighing, the GAS muscle was recombined with its hydrogel fragments (if any) and saved for either histological or biochemical analysis. Specifically, tissue from one limb was saved for biochemical analysis, while the other was preserved for histological analysis. For biochemical analysis, the muscle with any hydrogel fragments was flash-frozen in liquid nitrogen and stored at -80°C until further use. For histological analysis, the muscle with any histological fragments was embedded in optimal cutting temperature embedding medium (OCT, Thermo Fisher Scientific, Waltham, MA, USA, Catalog No. 23-730-571) in a 15×15×5 mm mold and then dipped in liquid nitrogen until the OCT completely solidified. The muscle samples were stored at -80°C until needed for further analysis.

### 2.7. Protein Expression Analysis

The GAS muscle along with any hydrogel fragments that was preserved for biochemical analysis (n=6/group) was homogenized on ice in 1× Pierce^TM^ radioimmunoprecipitation assay (RIPA) lysis buffer (Thermo Fisher Scientific, Waltham, MA, USA) containing a protease inhibitor cocktail (1:100; Sigma-Aldrich, a Millipore Sigma company, Saint Louis, MO, USA, Catalog No. P8340). The protein concentrations were quantified using a Pierce^TM^ Bicinchoninic Acid (BCA) Protein Assay kit (Thermo Fisher Scientific, Waltham, MA, USA, Catalog No. 23225). For each sample, equal amounts of protein (60 μg) were loaded on 4–20% precast polyacrylamide gels (Bio-Rad, Hercules, CA, USA, Catalog No. 4568096). The protein was resolved and separated based on its molecular weight by running a Sodium Dodecyl Sulfate Polyacrylamide Gel Electrophoresis (SDS-PAGE). The resolved protein bands were transferred onto nitrocellulose membranes and stained with Ponceau S (TOCRIS, Biotechne Corporation, Minneapolis, MN, USA Catalog No. 5225) to confirm equal protein loading. The protein bands were blocked for 1 hour at room temperature in Tris-buffered saline containing 0.05% (vol/vol) Tween 20 (TBST) and 5% (wt./vol) non-fat dried milk. Membranes were then incubated overnight at 4°C in TBST containing 5% bovine serum albumin (BSA, Sigma Aldrich, parent company Millipore Sigma, Saint Louis, MO, USA, Catalog No. 126575) and primary antibodies diluted at 1:1000. Primary antibodies for probing protein included MyoD (Invitrogen, Thermo Fisher Scientific, Waltham, MA, USA, Catalog No. MA1-41017), anti-myogenin (Sigma-Aldrich, parent company Millipore Sigma, Saint Louis, MO, USA, Catalog No. MAB3876), alpha-actinin (Cell Signaling Technology, Danvers, MA, Catalog No. 6487), desmin (Catalog No. ab15200), inducible nitric oxide synthase (iNOS, Catalog No. ab49999), and arginase (Catalog No. ab91279) (Abcam, Cambridge, MA, USA). After overnight incubation at 4°C, membranes were rinsed three times with TBST for 5 minutes and then incubated at room temperature for 1 hour in TBST containing 5% non-fat dried milk and the appropriate horseradish peroxidase-conjugated secondary antibody (1:500, Invitrogen, Thermo Fisher Scientific, Waltham, MA, USA). Membranes were rinsed three times with TBST for 5 minutes before exposure to chemiluminescent developer (Bio-Rad, Hercules, CA, USA, Catalog No. 1705061). The membranes were imaged using the Bio-Rad ChemiDoc Imaging System (BioRad, Hercules, CA, USA). Quantification of band intensity was performed using Image J. The data were normalized to GAPDH (Cell Signaling Technology, Danvers, MA, Catalog No. 5174) and used for quantitative analysis.

### 2.8. Histological Analysis

GAS muscle along with any hydrogel fragments that was preserved for histological analysis (n=6/group) were mounted for cryosectioning on stubs using OCT. Transverse cryosections (15 μm) were cut and mounted on charged gold microscope slides (ThermoFisher Scientific, Waltham, MA, USA, Catalog No. 151-188-48). Muscle cross-sections were stained with hematoxylin (Sigma-Aldrich, parent company Millipore Sigma, Saint Louis, MO, USA, Catalog No. MHS16) and eosin (Sigma-Aldrich, parent company Millipore Sigma, Saint Louis, MO, USA, Catalog No. HT110116) (H&E) to visualize the myofiber morphology. Histological analysis was performed on tissue cross-sections, which comprised the VML defect area, remaining muscle mass (RMM), as well as hydrogel fragments either integrated or non-integrated with the musculature.

For immunofluorescent staining, sections were fixed using ice-cold acetone or 4% paraformaldehyde, then permeabilized with a solution of 0.1% Triton X-100 in 1× PBS, and blocked in a solution containing 5% goat serum, 1% bovine serum albumin (BSA), and 0.05% Tween-20. The incubation buffer for both primary and secondary antibodies consisted of 1% BSA and 0.1% Triton X-100 in 1× PBS. The muscle sections were incubated overnight at 4 °C with primary antibodies diluted in incubation buffer. To determine macrophage infiltration, muscle cross-sections were stained with F4/80 (1:50, ThermoFisher Scientific, Waltham, MA, USA, Catalog No. MA5-16624), and the basal lamina of myofibers was identified with laminin antibody (1:200, ThermoFisher Scientific, Waltham, MA, Catalog No. PA1-16730). To identify vasculature, muscle cross sections were stained with von Willebrand factor antibody (vWF; 1:500, Abcam, Cambridge, MA, USA Catalog No. ab287967), and extracellular matrix deposition was determined using wheat germ agglutinin (WGA; 1:200, Thermo Fisher Scientific, Waltham, MA, USA, Catalog No. w11262). Appropriate fluorochrome-conjugated secondary antibodies for F4/80 (1:100, Invitrogen, parent company Thermo Fisher Scientific, Waltham, MA, USA, Catalog No. A32731), laminin (Invitrogen, parent company Thermo Fisher Scientific, Waltham, MA, USA, Catalog No. 41481582), and vWF (Abcam, Cambridge, MA, USA Catalog No. a11034) were diluted at 1:100. Sections were incubated with it at room temperature for 1 hour. Cell nuclei in muscle sections were stained and identified using 4’, 6-diamino-2-phenylindole (DAPI, Thermo Fisher Scientific, Waltham, MA, USA, Catalog No. 62248). Vectashield (Vector Laboratories, Newark, CA, USA, Catalog No. H-1000) was used as mounting media. All sections were cover-slipped, sealed with nail polish, and stored at 4°C for future analysis. The stained muscle sections were scanned using a Leica SP6 Epifluorescence microscope (Research Microscopy and Histology Core, SLU) to obtain 20X images.

WGA has been previously used in many studies to quantify fibrotic tissue deposition in skeletal [63, 64] and cardiac muscle [65]. For quantifying vWF and WGA stains, the defect and RMM, were outlined separately using ImageJ/FIJI and analyzed. The defect region was identified by the high density of nuclei (DAPI^+^) and low density of myofibers. The hydrogel embedded and integrated within the defect region was identified by its unique morphology and by nuclear deposition at the boundary, but with little to no nuclear infiltration within the hydrogel. The area of muscle tissue, excluding both defect and hydrogel, was identified as RMM. The non-integrated hydrogel fragment did not show positive staining for either vWF or WGA and was excluded from analysis. Muscle sections stained with vWF were used to quantify blood vessel percent area via AngioTool software (0.6a, National Cancer Institute, USA). The WGA stain on muscle sections was used to quantify the percent (%) area of fibrotic tissue deposition via thresholding using ImageJ/FIJI.

The muscle sections stained with laminin and F4/80 antibodies were used to quantify the cross-sectional area of myofibers and macrophage infiltration, respectively. Because macrophage infiltration (F4/80^+^) was primarily restricted to the defect region, a split analysis of defect and RMM was not performed. F4/80^+^ macrophages were present around and within both the integrated and the non-integrated hydrogel fragments. Thus, both hydrogel fragments were included in the analysis. The amount of macrophage infiltration was quantified using thresholding in Image J. The MyoQuant image analysis program [66] was used to quantify the total number of laminin^+^ myofibers and their cross-sectional area (CSA). The area of myofibers, obtained in μm², was segregated into size bins in the range of <500 >6000 μm². Myofibers (laminin+) with centrally located nuclei (DAPI^+^) were manually counted using the multi-point tool in ImageJ/FIJI.

### 2.9. Protein Cargo Analysis of EVs

The biological signals carried by EVs as cytokines, chemokines, and growth factors, were analyzed using Proteome Profiler Mouse XL Cytokine Array (R&D Systems, Biotechne Corporation, Minneapolis, MN, USA, Catalog No. ARY028). Norm-EVs and Hypo-EVs were pooled from three independent isolations and were processed on separate membranes. EVs were resuspended in 1× sterile PBS at a concentration of 100 μg/ml and lysed using a 10× Lysis Buffer (from the kit for Exo-Check^TM^ Exosome Antibody Array, System Biosciences, Palo Alto, CA, USA, Catalog No. EXORAY200B-4), which was diluted to 1× directly into the EV solution to dissociate the EVs. Based on the manufacturer’s instructions, the membranes were blocked for 1 hour at room temperature, then incubated overnight at 4 °C with the lysed EV sample. After washing, the membranes were incubated with detection antibody cocktail for 1 hour at room temperature, washed again, and subsequently incubated with streptavidin-HRP for 30 minutes at room temperature. The membranes were washed again, and the chemiluminescent signals were visualized using BioRad ChemiDoc Imaging System (BioRad, Hercules, CA, US). The spot intensities were quantified using ImageJ software, and duplicate spots were averaged. The values were normalized to the positive control spots on each membrane. The fold changes between Norm-EVs and Hypo-EVs were calculated as log2 and plotted.

### 2.10. Statistical Analysis

The data is presented as the mean ± standard error of the mean (SEM), with statistical significance set at p < 0.05. Statistical analysis and graph generation were conducted using GraphPad Prism 10. A one-way analysis of variance (ANOVA) with Fisher’s least significant difference (LSD) post-hoc test was performed to detect statistical significance. Fiber-size distribution data were evaluated using a two-way ANOVA. The EV size mean, mode, and span were statistically assessed using the Mann-Whitney U test.

## 3. Results

### 3.1. Hypoxic and normoxic EVs differ in morphology and surface marker expression

The size and concentration of Norm-EVs and Hypo-EVs were determined using NTA (Fig. 1A). Hypo-EVs had a mean concentration of 1.69E+07 particles/ml whereas Norm-EVs had a concentration of 1.12E+07 particles/ml. Thus, culturing BM-MSCs in hypoxia (3% O_2_) resulted in a 1.5-fold higher production of EVs (Fig. 1B, Mann Whitney test, p=0.0002). There were no significant differences between Hypo-EVs and Norm-EVs in the mean particle size (Fig. 1C, Mann Whitney test, p>0.999). The size span of Norm-EVs was significantly lower than Hypo-EVs, indicating that Norm-EVs were more homogenous in size than Hypo-EVs (Fig. 1D, Mann-Whitney test, p=0.0317). Even though normoxic-preconditioning of MSCs produced EVs at lower concentrations, the Norm-EVs had higher expression of several characteristic EV markers, such as FLOT 1, ICAM, CD81, CD63, and ANXA5, compared to Hypo-EVs. Both Norm-EVs and Hypo-EVs were negative for GM130, a marker of cellular contamination [67, 68], thus indicating an EV population free from cellular debris (Fig. 1E-G).

**Figure 1:**
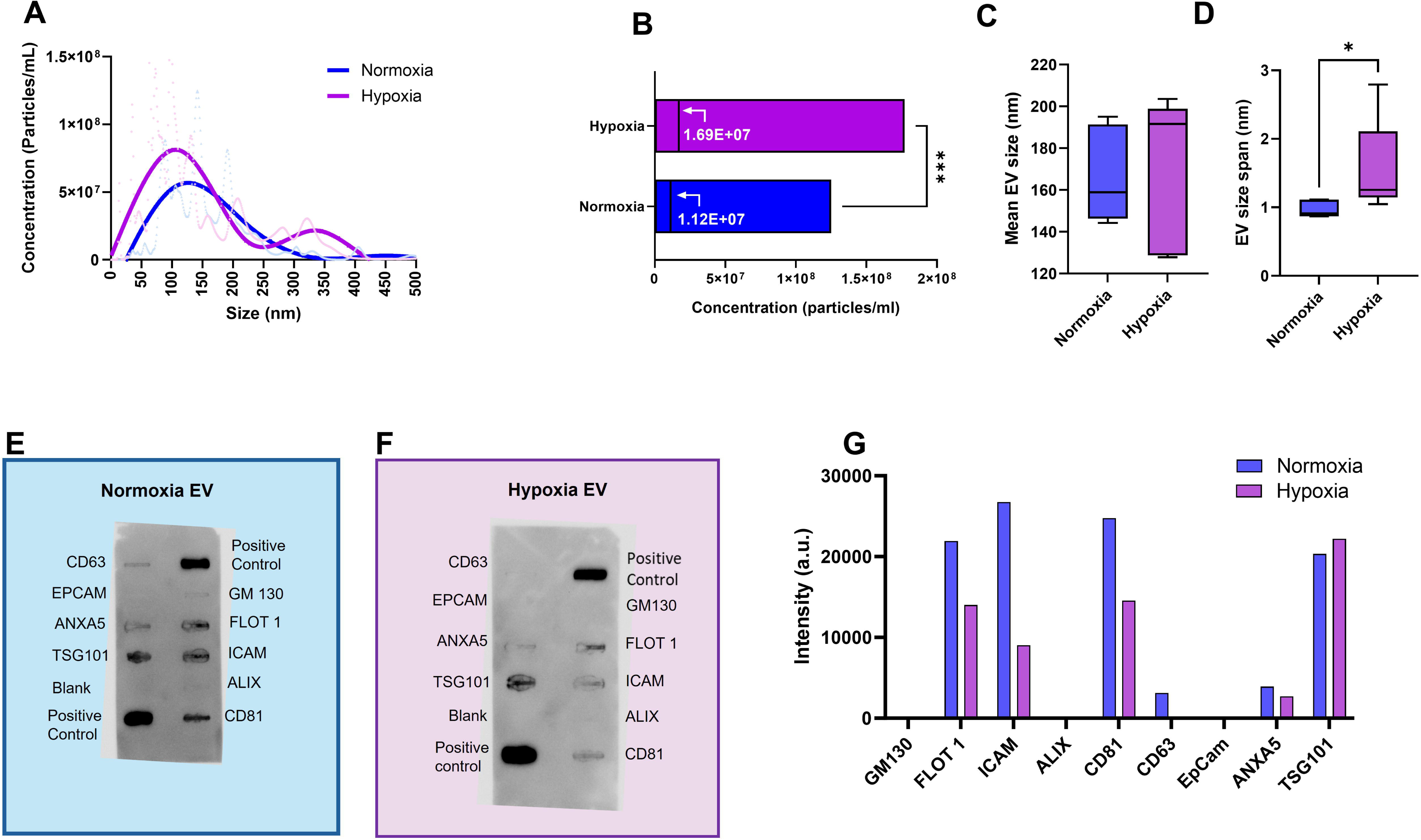
Nanoparticle Tracking Analysis (NTA) showed higher EV concentration (particles/mL) from MSCs cultured under hypoxia compared to normoxia. (A) The size distribution of EVs is displayed. (B) The average concentrations are shown for both groups. (C) No significant differences were observed in the mean size of hypoxia or normoxia derived EVs (D) Hypoxia-derived EVs had a significantly larger size span compared to normoxia derived EVs. Exo Check^TM^ Exosome Antibody Array depicting expression of characteristic EV markers by (E) Normoxia EVs and (F) Hypoxia EVs is shown. (G) Hypoxia EVs had lower expression for several markers (e.g., FLOT 1, ICAM, CD81, and ANXA5) relative to Normoxia EVs. The Mann-Whitney test was used for statistical analysis. “*” indicates a significant difference (*p<0.05, ***p<0.001, ****p<0.0001) between groups.

### 3.2. Immunomodulatory effect of hypoxic and normoxic EVs *in vitro*

LPS-stimulated macrophages (MΦs) were cultured in media alone (control) or with media containing Norm-EVs or Hypo-EVs. MΦ culture media were used as a control. To assess the effect of Norm-EVs and Hypo-EVs on MΦ polarization *in vitro*, production of pro-inflammatory M-1 phenotype-associated cytokines IL-6 and TNF-α, and anti-inflammatory M2 phenotype-associated cytokine, VEGF, by MΦs was quantified. Norm-EVs promoted higher IL-6 production than media (Figure 2A, ANOVA p= 0.0047) or Hypo-EVs (p= 0.0358) (Figure 2A, ANOVA p= 0.0130). Quantification of TNF-α (Figure 2B, ANOVA p=0.0209) revealed that both Norm-EVs and Hypo-EVs suppressed its production compared to media (media vs. Norm-EV, p=0.0248, media vs Hypo-EV, p=0.0093). While both Hypo-EVs and Norm-EVs increased production of VEGF from MΦs compared to media alone (Figure 2C, ANOVA p=0.007, plain media vs Hypo-EVs, p=0.0024, plain media vs Norm-EV, p=0.0003), VEGF production was increased more in response to Norm-EVs than to Hypo-EVs (p=0.0396). Overall, lower production of IL-6 and TNF-α, and increased production of VEGF, in response to Hypo-EVs indicate that Hypo-EVs deliver signals that polarize MΦs towards the anti-inflammatory M2 phenotype. In comparison, Norm-EVs promote IL-6 and VEGF production but suppress TNF-α production, indicating that they deliver cues that can polarize MΦs into both M1 and M2 phenotypes.

**Figure 2:**
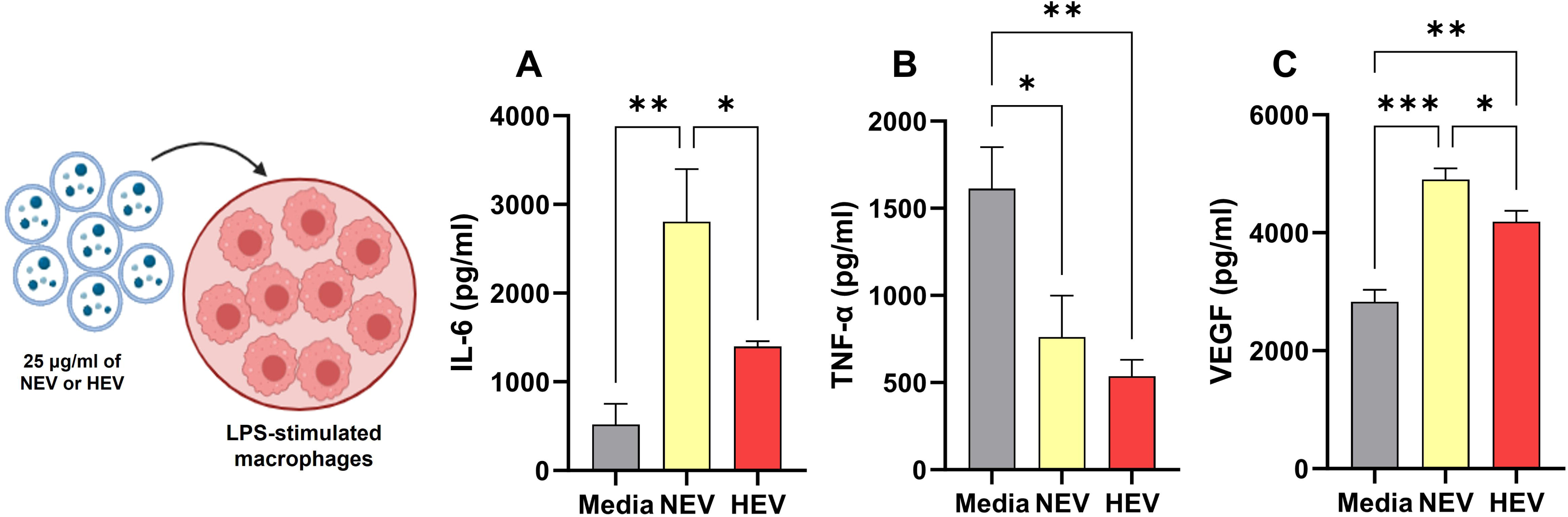
The release of (A) Interleukin-6 (IL-6), (B) Tumor Necrosis Factor-α (TNF-α), and (C) Vascular Endothelial Factor (VEGF) released by lipopolysaccharide (LPS) stimulated macrophages in response to plain media, Norm-EVs, or Hypo-EVs was quantified. Statistical analysis was performed using One-Way ANOVA. “*” indicates a significant difference (*p<0.05, ***p<0.001, ****p<0.0001) between groups.

### 3.3. Release kinetics of EVs from fibrin hydrogels

The release of EVs from fibrin hydrogels was studied using Exo-Check^TM^ Exosome Antibody Array. The PBS collected from EV-containing hydrogels on day 1 (Figure 3A) and day 2 (Figure 3B) showed positive expression for several EV markers, including ICAM, CD63, TSG101, ANXA5, and EpCAM (Figure 3E). However, PBS collected on day 3 (Figure 3C) showed no positive expression for any EV markers. Unexpectedly, PBS collected from the plain fibrin hydrogel (control) on day 1 also showed positive expression of specific EV markers (Figure 3D). To account for this, the expression level of EV markers by EV containing fibrin gels on days 1 and 2 was normalized to the expression level of EV markers by plain fibrin gel (control). The fold change values were calculated relative to control, which is indicated by a dotted line in the bar graph (Figure 3E).

**Figure 3:**
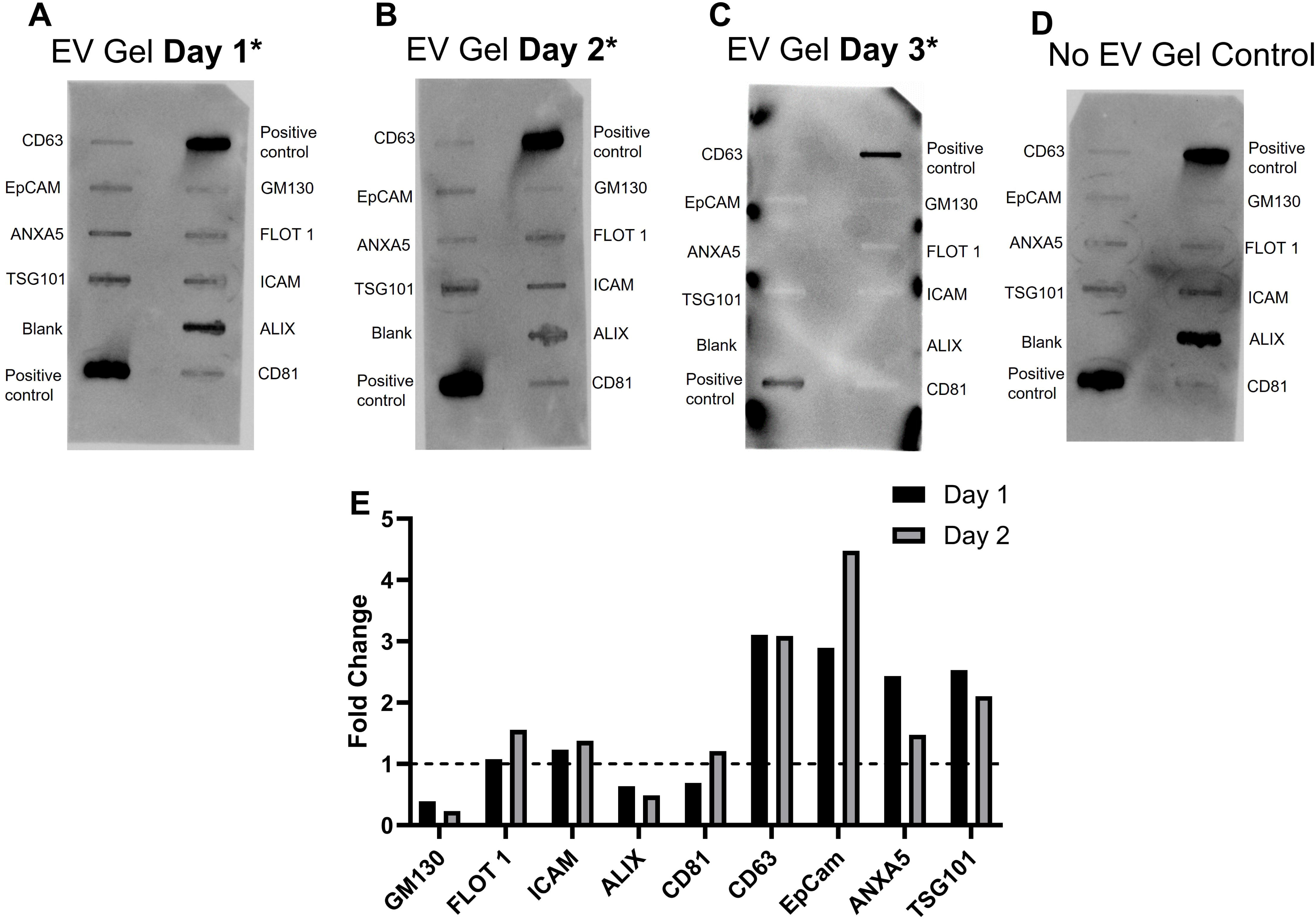
The release kinetics of EVs from fibrin hydrogels was determined using Exo-Check^TM^ Exosome Antibody Array on (A) Day 1, (B) Day 2, and (C) Day 3. (D) A control fibrin hydrogel (without EVs) was also tested for characteristic EV markers. (E) Quantification of band intensity is shown. Data was normalized to control gel values (dotted black lines). EV encapsulated hydrogels release EVs on days 1 and 2 but not on day 3.

### 3.4. Therapeutic effects of EVs on muscle recovery post-injury

The biopsy mass removed from the gastrocnemius-soleus complex to create the VML injury was weighed, and no significant differences were observed among the experimental groups (Figure 4A, Kruskal-Wallis, p = 0.9485). This indicates that the VML defect created was consistent across the treatment groups. On day 14 post-injury, muscle harvested from mice that received the fibrin hydrogel encapsulated Norm-EVs had significantly higher muscle mass (Figure 4B, one-way ANOVA, p = 0.0452, Norm-EVs vs. Hypo-EVs, p= 0.0132) compared to mice that received fibrin hydrogel encapsulated Hypo-EVs.

**Figure 4:**
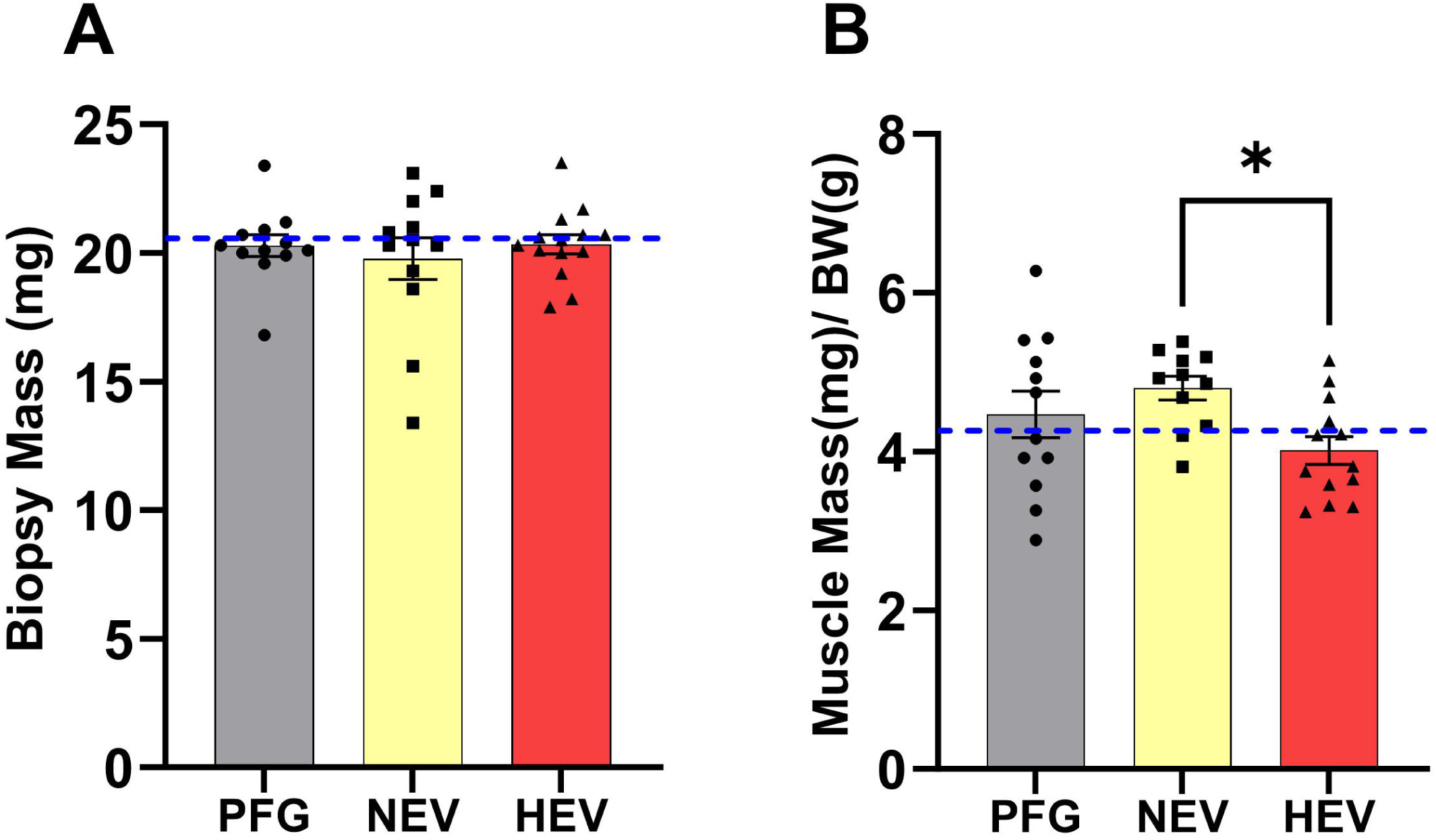
(A) The biopsy mass removed during surgery to create VML defect was similar between the experimental groups. Muscle harvested from both legs at a 14-day recovery time point was weighed and normalized to body weight. (B) Norm-EV treatment resulted in greater muscle mass than Hypo-EV treatment. Statistical analysis was performed using one-way ANOVA. “*” indicates a significant statistical difference (*p<0.05).

### 3.5. Myogenesis and muscle repair

Protein lysates obtained from muscle tissue were tested for expression of several myogenic and immunogenic markers. Representative immunoblots are shown Supplementary Fig. 1 A and B. There were no significant differences observed among experimental groups in the expression of early myogenic markers; myoblast determination protein (MyoD; one-way ANOVA, p=0.5145), and myogenin (one-way ANOVA, p=0.4007) (Supplementary Fig. 1 C and D), late myogenic markers; desmin (one-way ANOVA, p=0.0823) and alpha-actinin (one-way ANOVA, p=0.1793) (Supplementary Fig. 1 E and F), pro-inflammatory marker; iNOS (one-way ANOVA, p=0.7628) (Supplementary Fig. 1G) and anti-inflammatory marker; arginase (one-way ANOVA, p=0.6379) (Supplementary Fig. 1H).

Muscle cross-sections were stained with laminin and DAPI on day 14 post-injury (Figure 5A). There were no significant differences observed in the number of myofibers (Figure 5B, one-way ANOVA, p=0.6576) and the mean cross-sectional area of myofibers (Figure 5C, one-way ANOVA, p=0.9461) across the experimental groups. There was a significantly higher number of myofibers (Figure 5D, one-way ANOVA, p=0.0378) with centrally located nuclei (CLN) in the fibrin hydrogel encapsulated Norm-EV treatment (p=0.0440) and plain fibrin gel treatment (p=0.0176) compared to fibrin hydrogel encapsulated Hypo-EV treatment. Quantitative analysis of percentage of myofibers in different area size bins ranging from <500 μm^2^ to >6000 μm^2^ (Figure 5E, two-way ANOVA, interaction, p=0.0477, row factor, p=<0.0001) showed that fibrin hydrogel encapsulated Norm-EV treatment resulted in increased number of small-sized fibers (<500 μm^2^) relative to plain fibrin gel (p= 0.0003) or fibrin hydrogel encapsulated Hypo-EV treatment (p=0.0019). Plain fibrin gel treatment resulted in the formation of higher numbers of myofibers in the size range of 500-999 μm² relative to fibrin hydrogel-encapsulated Norm-EV treatment (p = 0.0162). Thus, these results indicate that fibrin hydrogel-encapsulated Norm-EVs are shifting the myofiber size distribution to the left, due to a higher percentage of small-sized myofibers.

**Figure 5:**
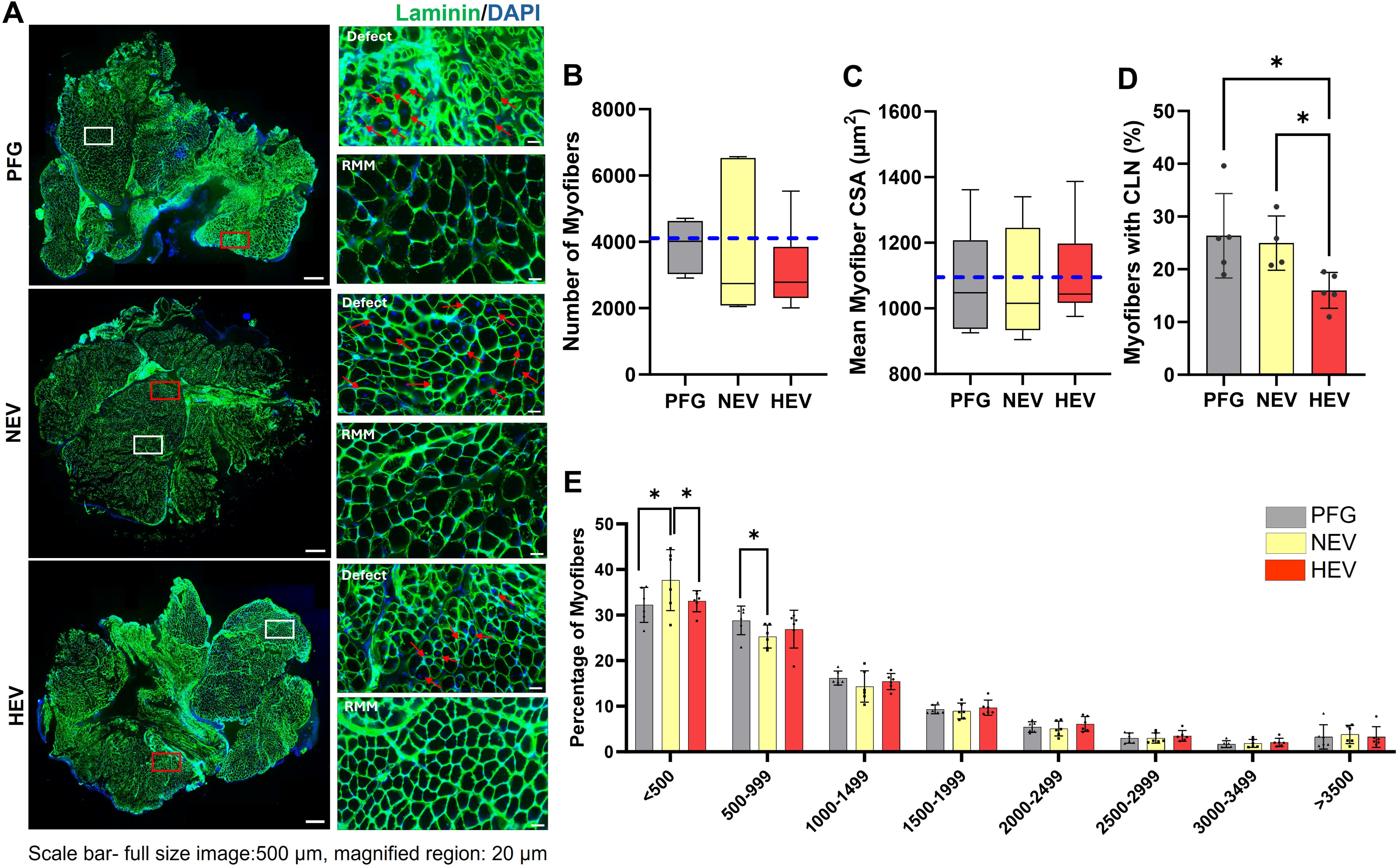
(A) Myofibers were stained with laminin and DAPI. The regions adjacent to the defect (red box) and remaining muscle mass (RMM; white box) are magnified. Myofibers with centrally located nuclei (CLN) are identified with red arrows. The (A) number of myofibers, (B) mean myofiber cross-sectional area (CSA), and (D) percentage of myofibers with CLN were quantified. (E) The size distribution of myofibers is shown. The dashed line in graphs (B) and (C) represents the average value obtained for n=2 VML group samples. Scale bar for full-size images is 500 μm, and magnified images are 20 μm. Statistical analysis was performed using One-Way or Two-way ANOVA. “*” indicates a significant statistical difference (**p<0.01, ***p<0.001, ****p<0.0001)

### 3.6. Macrophage infiltration, angiogenesis, and fibrosis

Macrophage (F4/80^+^) infiltration in response to plain fibrin gel, Norm-EV, or Hypo-EV encapsulated fibrin gel was quantified (Figure 6A). F4/80^+^ macrophages were detected in all three groups primarily in the defect area but also around both integrated and non-integrated hydrogel fragments (Figure 6B, one-way ANOVA, p=0.0150). Fibrin gel encapsulated with Norm-EVs (one-way ANOVA, p= 0.0058) or Hypo-EVs (one-way ANOVA, p=0.0272) significantly suppressed macrophage infiltration in the defect area compared to plain fibrin gel treatment. The integrated hydrogel fragment is shown with yellow dashed lines. The non-integrated hydrogel fragment is not shown.

**Figure 6:**
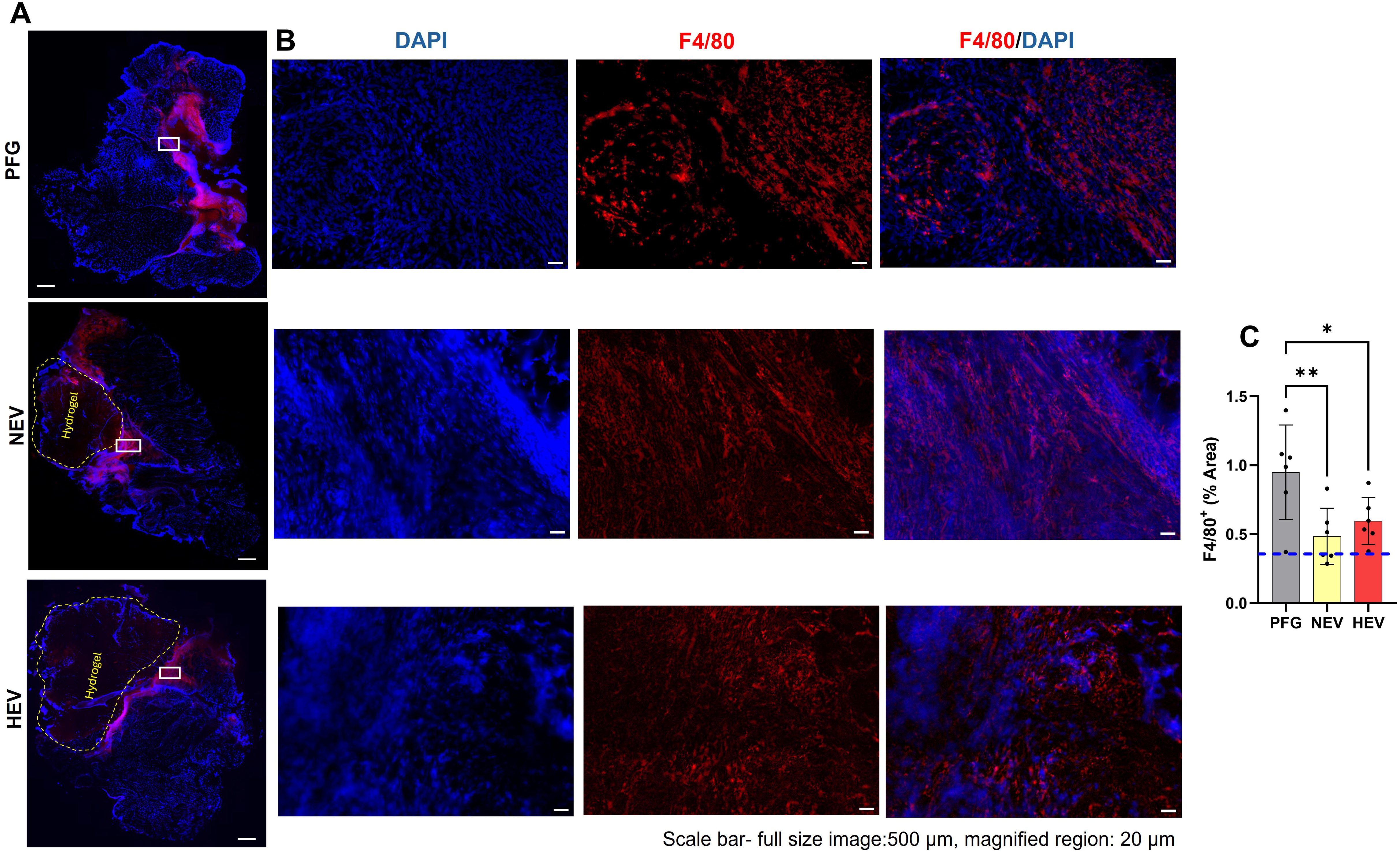
(A) Muscle cross-sections were immunostained with F4/80 and DAPI to identify macrophage infiltration and cell nuclei, respectively. The yellow dashed lines indicate the integrated hydrogel fragment. The white box outlines a dense F4/80+ region, which is magnified. (B) The magnified images highlight the punctate nature of F4/80 staining as well as its co-localization with DAPI. Scale bar for full-size images is 500 μm, and magnified images are 20 μm. (C) Quantification of F4/80+ stained area showed that both Norm-EV and Hypo-EV treatment reduced macrophage infiltration in the defect region relative to PFG. Statistical analysis was performed using one-way ANOVA. “*” indicates a significant statistical difference (**p<0.01, ***p<0.001, ****p<0.0001).

Blood vessel (vWF^+^) percentage area was quantified in the defect area, remaining muscle mass (RMM) area, and entire muscle cross-section area (i.e., total) on day 14 post-injury (Figure 7A). Only the integrated hydrogels showed vWF^+^ blood vessels. Fibrin hydrogel encapsulated with Norm-EVs resulted in a significantly higher density of vWF+ blood vessels (Figure 7B, one-way ANOVA, p=0.0295) in the defect region compared to fibrin hydrogel encapsulated with Hypo-EVs (one-way ANOVA, p=0.0178) or PFG (one-way ANOVA, p=0.0226). There were no significant differences observed in blood vessel percentage area in the RMM area (Figure 7C, one-way ANOVA, p=0.5419) and total muscle tissue area cross-section (Figure 7D, one-way ANOVA, p=0.0662).

**Figure 7:**
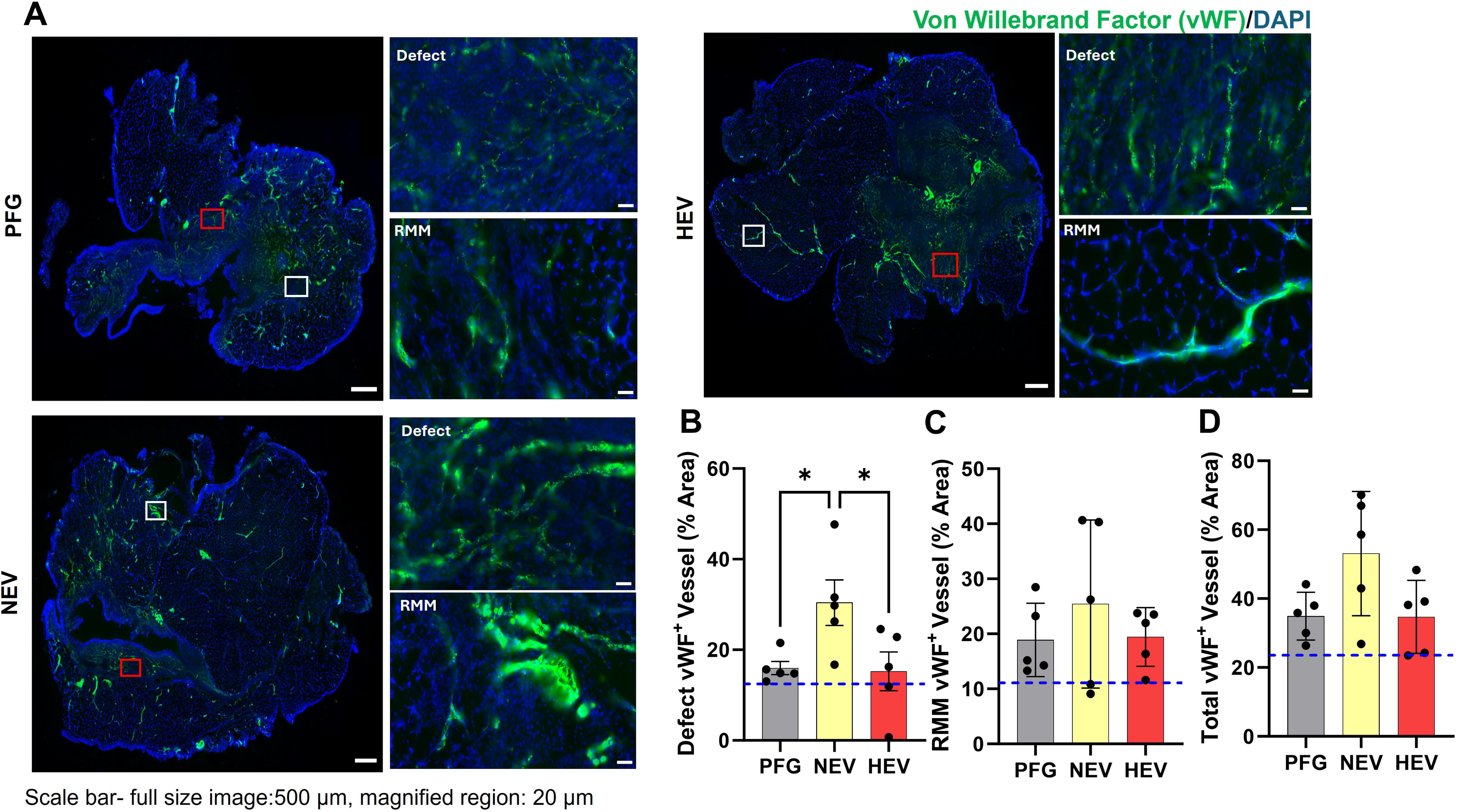
(A) Muscle cross sections were immunostained with von Willebrand factor (vWF) and DAPI to identify blood capillaries and cell nuclei respectively. Red box outlines the blood vessels in the defect region, and the white box outlines the blood vessels in the RMM area. Scale bar for full size section: 500 μm, scale bar for zoomed in section: 20 μm. (B) Norm-EV treatment promoted formation of vWF+ blood vessels in the defect region compared to Hypo-EV treatment or PFG. No significant difference was observed in the area percentage of vWF+ stained blood vessels in (C) in the RMM region and (D) and total muscle cross-section. Dashed line in graph (B), (C) and (D) represents the average value obtained for n=2 VML group samples. Statistical analysis was performed using One-way ANOVA. “*” indicates a significant statistical difference (**p<0.01, ***p<0.001, ****p<0.0001).

Muscle cross-sections were stained with WGA to assess fibrotic tissue deposition (Figure 8A). Only the integrated hydrogel fragments stained positive for WGA. No significant differences were observed among experimental groups in terms of fibrotic tissue deposition in the defect area (Figure 8B, one-way ANOVA, p=0.7700), in the RMM area (Figure 8C, one-way ANOVA, p=0.2164), entire muscle cross sections (Figure 8D, one-way ANOVA, p=0.7994).

**Figure 8:**
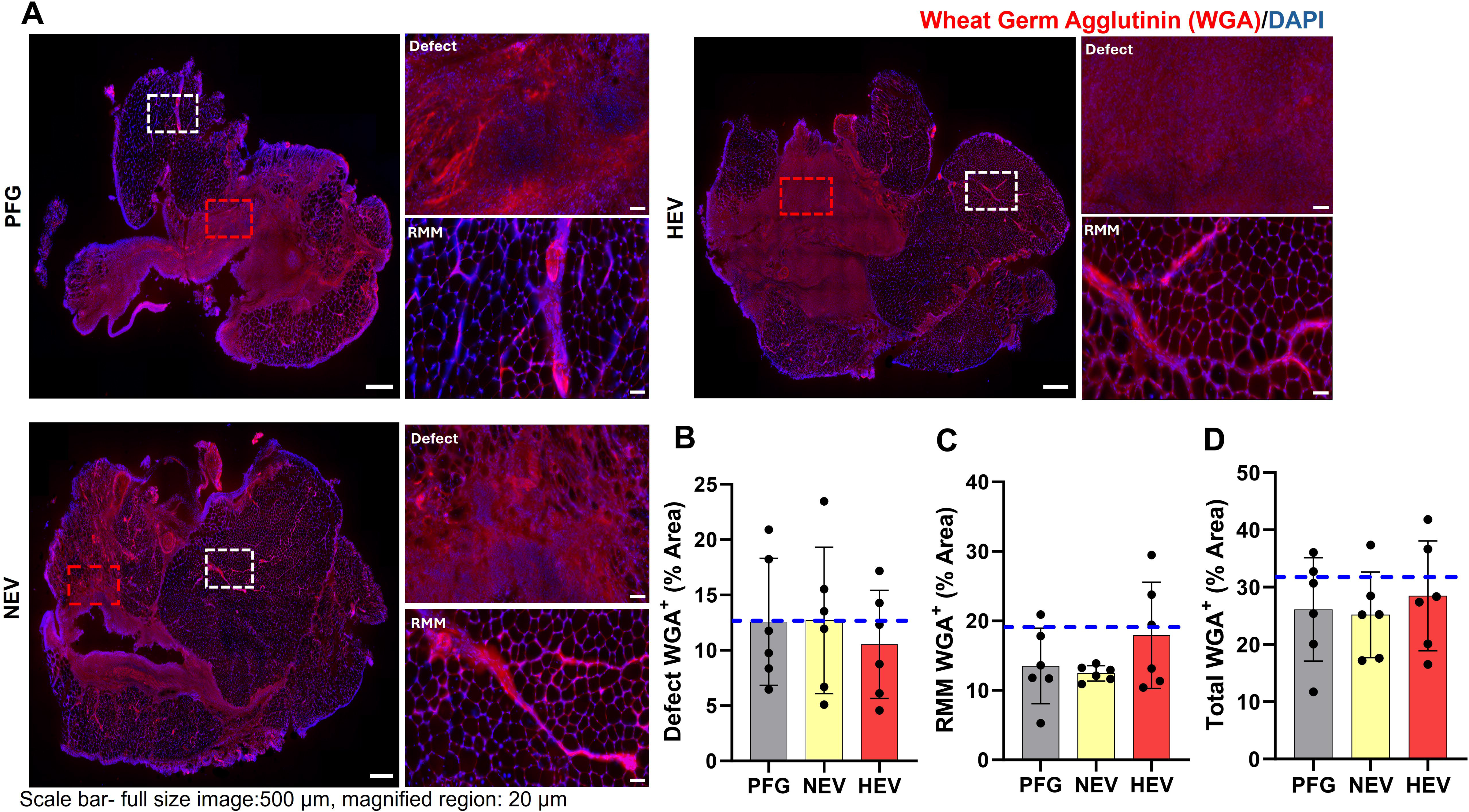
(A) Muscle cross-sections were immunostained with Wheat Germ Agglutinin (WGA) to identify extracellular matrix (ECM) deposition. The defect region (red box) and remaining muscle mass (RMM; white box) are magnified. Scale bar for full-size images is 500 _μ_m, and magnified images are 20 _μ_m. There were no significant differences in ECM deposition in (B) defect region, (C) RMM, and (D) full muscle section. Statistical analysis was performed using one-way ANOVA.

The non-integrated hydrogel fragments weighed upon tissue harvest are graphed in Supplementary Fig. 2. The non-integrated hydrogel fragments weighed significantly higher in the PFG group relative to both the Norm-EV and Hypo-EV groups (one-way ANOVA, p = 0.0014). It should be noted, however, that in the case of EV-containing hydrogels, a higher percentage of injured muscles (∼83%) retained measurable hydrogel pieces, despite the hydrogels having lower overall mass upon harvest (PFG vs. NEV, one-way ANOVA, p = 0.0028 and PFG vs. HEV, p=0.005). These results suggest improved retention and integration of the EV-containing hydrogels at the injury site, as well as consistent, gradual degradation and remodeling of the hydrogels. Conversely, PFG showed lower retention rates (∼50%) and higher mass when harvested, suggesting lower integration with the surrounding musculature, higher displacement rates, and inconsistent degradation and remodeling.

### 3.7 EV cargo profiling

To identify and differentiate the biological cargo carried by Norm-EVs and Hypo-EVs, we examined the expression of 111 mouse proteins using cytokine array (Fig. 9A -B). The upregulated (log2 fold changes>0.5) and downregulated proteins (log2 fold changes <0.5) were highlighted using colored boxes on the array and plotted on a heat map (Fig. 9C). Norm-EVs showed higher expression of certain interleukins such as IL-33[69], chemokines, such as C-C motif chemokine ligand 5 (CCL5), also known as regulated upon activation, normal T-cell expressed and secreted (RANTES), C-X-C motif chemokine ligand 3 (CXCL3), also known as Fractalkine, CXCL2, also known as macrophage infiltration protein (MIP)-2; tumor necrosis factor receptor superfamily (TNFRSF) cytokines such CD40/TNFRSF5[69], Osteoprotogerin /TNFRSF11B[70]; cytokines, such as Interferon (IFN)-γ, insulin-like growth factor binding protein 6 (IGFBP-6), VCAM ( vascular cell adhesion protein)/CD106, Leptin[69], LIF[71]; glycoproteins such as Endostatin[72], Fetuin A/AHSG (alpha-2-Heremans-Schmid glycoprotein), [73], myeloperoxidase[74]; low-density lipoprotein receptor (LDL-R) [75]; non-glycosylated proteins such as Cystatin-C[76]; peptides such as regenerative islet-derived 3 gamma (Reg3G) [77]; matricellular protein such as Osteopontin[78], WNT1-inducible signaling pathway protein-1 (WISP-1), also known as cellular communication network factor 4 (CCN4) [79]; and growth factors such as Coagulation Factor III/Tissue Factor and macrophage colony stimulating factor (M-CSF) [80]. Only one cytokine/adipokine, adiponectin, also known as adipocyte complement-related protein 30 (Acrp30) [81] was downregulated in Norm-EVs compared to Hypo-EVs.

**Figure 9:**
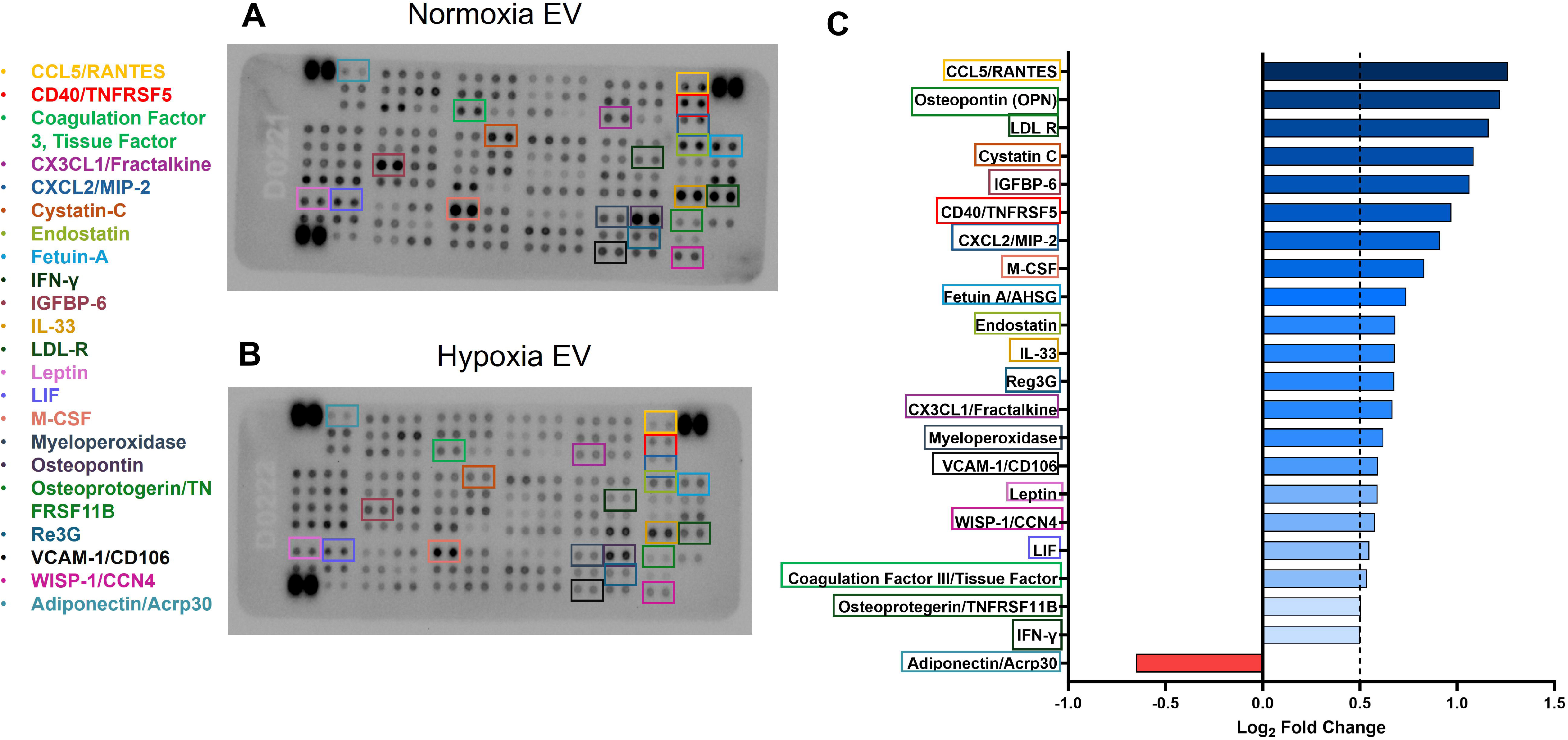
The levels of proteins within both (A) Norm-EVs and (B) Hypo-EVs were quantified using a protein array. The proteins of interest are labeled with colored boxes on both membranes. (C) The comparison between Norm-EV and Hypo-EV is represented as log2 fold changes. Blue color intensity indicates increased expression, while the red color indicates decreased expression.

## 4. Discussion

This study compared and evaluated the regenerative and immunomodulatory potential of normoxic vs hypoxic pre-conditioned BM-MSC-derived EVs (Norm-EV vs. Hypo-EV) in a murine model of VML. We demonstrate that while cyclic hypoxia (3% O_2_) enhances EV yield, it reduces the regenerative potential of EVs. In contrast, culturing MSCs in normoxia (21% O_2_) produces EVs that promote muscle regeneration and angiogenesis. These findings reveal that increased EV production does not compensate for loss in regenerative potency, highlighting the importance of EV quality over quantity for VML therapies.

Previous studies have shown the therapeutic efficacy of MSC-EVs in promoting skeletal muscle regeneration following both acute and chronic injuries [34, 35, 82, 83]. Our study specifically investigated whether hypoxic or normoxic culture conditions alter MSC-EV functionality in a VML model. Our results show that Norm-EVs increase the percentage of small-sized (<500 µm²) myofibers. Moreover, both PFG and Norm-EV treatments significantly increased the presence of myofibers with centrally located nuclei (CLN) compared with the Hypo-EV treatment. In muscle repair or regeneration, smaller myofibers are often indicative of newly formed or immature myofibers that emerge during the process of myogenesis [84, 85].

This process involves the activation, proliferation, and differentiation of muscle stem cells (satellite cells) into myoblasts, which then fuse to form new myofibers or repair damaged ones [86]. The increased percentage of myofibers with CLN alongside a higher frequency of small-diameter myofibers (<500 µm^2^), indicates elevated myogenesis in response to Norm-EV treatment. Although we did not observe any changes in protein expression of myogenic markers, such as MyoD and myogenin, all treatment groups showed ∼3-fold higher expression than the untreated VML-injured muscles, indicating a heightened myogenic response which was not further elevated with EV incorporation.

Hypoxic preconditioning has been shown to result in pro-angiogenic effects on MSCs [39, 87]. However, we observed higher capillary density following Norm-EV treatment compared to Hypo-EV treatment in VML-injured muscles. A recent study by Sicco *et al* compared the angiogenic potential of EVs isolated from adipose tissue derived-MSCs maintained for 48 hours in normoxic (20% O_2_) vs hypoxic (1% O_2_) conditions using a Matrigel plug assay. Both normoxic and hypoxic MSC-derived EVs increased vessel density and the expression of angiogenic molecules, such as VEGF-A [88]. In another study, EVs were isolated from adipose tissue derived-MSCs cultured for 6 days under normoxic (21% O_2_) or hypoxic (5% O_2_) conditions. Human umbilical vein endothelial cells (HUVECs) were treated with the isolated EVs or EV-depleted supernatants from these normoxic or hypoxic MSCs. *In vitro*, hypoxic MSC-derived EVs significantly enhanced the angiogenic properties (vascular tube formation, branch points, and loops in vessels) of HUVECs [89]. In our study, we compared the effect of normoxic vs. hypoxic pre-conditioning of BM-MSCs on the angiogenic potential of derived EVs *in vivo*. Norm-EVs delivered via a fibrin hydrogel promoted greater blood vessel growth in the defect area than Hypo-EVs or PFG. The difference in the angiogenic potential of Hypo-EVs observed in our findings could be attributed to the tissue source of MSCs (bone marrow vs. adipose), as well as to the intensity (3% vs. 1 or 5%), duration (i.e., hours of hypoxia exposure), and type (continuous vs. cyclic) of hypoxia.

While Hypo-EVs lacked the myogenic and angiogenic potential of Norm-EVs, both EVs suppressed macrophage infiltration in the VML defect area. Macrophages also surrounded the hydrogel boundary but did not infiltrate the hydrogel. Our *in vitro* studies demonstrated that both Norm-EVs and Hypo-EVs interact with LPS-stimulated macrophages to modulate their secretion of M1- or M2-like trophic factors. M1-phenotype-associated macrophages produce pro-inflammatory cytokines such as IL-6, TNF-α, and IL-1β, which signal the clearance of necrotic cell debris and activate tissue-resident muscle stem cells/satellite cells. Conversely, M2-phenotype-associated macrophages are pro-reparative and secrete IL-4, IL-10, IGF-1, and VEGF, which reduce inflammation and promote the proliferation and differentiation of activated satellite cells [90]. Maintaining an optimal balance between the presence of M1 and M2 macrophages at the injury site is crucial to wound healing [91]. MSC-EVs derived under hypoxic and normoxic conditions have been shown to maintain M1/M2 balance *in vivo* in a mouse model of acute cardiotoxin-induced skeletal muscle injury [88]. Our *in vitro* studies demonstrated that in the presence of Hypo-EVs, macrophages increase the secretion of M2-associated factors, such as VEGF, but reduce the secretion of both IL-6 and TNF-α. On the other hand, macrophages incubated with Norm-EVs produce high amounts of IL-6 and VEGF, but lower amounts of TNF-α, suggesting the induction of a mixed M1/M2 phenotype. However, we did not observe any differences in the protein expression of iNOS (an M1-associated marker or arginase (an M2-associated marker) in the VML-injured muscles in response to PFG, Hypo-EV, or Norm-EV treatment. These results suggest that while both Hypo-EVs and Norm-EVs suppressed macrophage infiltration into the defect, they did not modulate the macrophage phenotype on day 14 post-injury. It is also possible that transient phenotypic changes occurred earlier (e.g., day 7 post-injury) and converged to a similar state by day 14. Because this study used a single time point for tissue analysis, transient changes in macrophage phenotype may have been missed. A comprehensive temporal analysis with multiple M1/M2 markers is necessary to substantiate these claims.

The extent of fibrin hydrogel remodeling mirrors the degree of macrophage infiltration into injured muscles. EV encapsulating fibrin hydrogels reduced macrophage infiltration and likely underwent slower degradation primarily through hydrolysis. In contrast, plain fibrin hydrogels elicited heightened macrophage activity and underwent accelerated degradation likely through both hydrolytic and enzymatic degradation.

To prolong the presence of EVs at the VML injury site, we encapsulated them within fibrin hydrogels. Studies suggest that EVs can bind to ECM proteins via several different mechanisms. Wang *et al.* performed a proteomic analysis of MSC-EVs and found several signaling and adhesion molecules in quantifiable amounts. MSC-EVs express integrin that can bind to ECM protein ligands such as fibronectin, collagen, and laminin. MSC EVs also express proteins such as CD44, CD47, vinculin, and cadherins that mediate cell adhesion [92]. Fibronectin and collagen are also rich in positively charged basic amino acids that can bind to negatively charged phosphates on EVs via hydrogen bonding [93]. Studies reported that thrombin cleaves fibrinogen, thereby exposing heparin-binding domains. EVs derived from human BM-derived MSCs are rich in heparin sulphate proteoglycans and can bind to heparin-binding domains on fibrin via electrostatic interactions [94, 95]. While the exact mechanism of interaction between fibrin and EVs is not well known, several studies have utilized fibrin hydrogels to deliver EVs [53–56]. These studies reported that a fibrin hydrogel helped prolong the presence of EVs at the injury site, resulting in better therapeutic outcomes than direct injection of EVs. We studied the release kinetics of EVs from the EV-encapsulated fibrin hydrogel *in vitro*. We detected EV release in the surrounding PBS for up to 48 hours, but our experiment did not account for EVs entrapped in the fibrin-thrombin network, which will be released as the hydrogel degrades. A plain fibrin hydrogel was used as a control, and the surrounding PBS also showed expression of specific EV markers. The fibrinogen used to fabricate these hydrogels was extracted from bovine plasma by the manufacturing company. The plasma can have circulating EVs or EVs that endogenously express fibrinogen on their surface, which can be easily co-precipitated during fibrinogen extraction [96, 97]. Furthermore, although we didn’t quantify the EVs encapsulated within the fibrin hydrogel, our *in vivo* results suggest that EVs were delivered to the injury site and had physiological effects on muscle recovery, angiogenesis, and immune modulation on day 14 post-injury.

The cytokine array confirmed that Norm-EVs carry a rich and diverse cargo with a complex proteome profile. This profile comprises various proteins involved in the regulation of angiogenesis (e.g., VCAM-1 [98], Endostatin [99]), myogenesis (e.g., IGFBP-6 [100, 101], Osteopontin [78]), neuromuscular communication (e.g., Reg3G [102]), immunomodulation/chemoattraction (e.g., IL-33 [103], CCL5 [104], CXCL2 [105]), metabolism (e.g., Fetuin-A [106], LDLR [107]). In contrast, Hypo-EVs showed a biologically quiescent proteome profile with markedly lower levels of almost all proteins. Future studies will perform a detailed comparison of the EV cargo from normoxic vs. hypoxic pre-conditioned MSCs. This study also highlights the need for studying other methods of MSC preconditioning, such as 3-D culture technology (e.g., cell spheroids, bioreactors, quantum cell expansion systems etc.) [108], electrical stimulation [109], chemical [110] or cytokine [108] mediated stimulation to boost both EV production and their therapeutic potential.

Our results also show that hypoxic pre-conditioning of BM-MSCs improved the production of EVs but altered their composition. The reduced expression of characteristic EV markers, such as FLOT1, ICAM, CD81, CD63, and ANAX5, in Hypo-EVs may have lowered their therapeutic effects. Future studies will investigate different oxygen tensions and hypoxia durations to optimize cargo packaging in EVs to promote skeletal muscle regeneration. It is plausible that hypoxia forces cells into survival mode, thereby disrupting normal functions such as protein synthesis and cargo packaging into EVs [111, 112]. Under stress, cells prioritize survival mechanisms over repair-promoting activities, which could make hypoxia EVs less equipped with key factors that are crucial for tissue repair [113, 114].

These findings are interesting and insightful, but the study has some limitations. For instance, we did not evaluate functional recovery in response to Hypo-EV and Norm-EV treatment post-VML, due to the single time-point used for analysis. Another limitation is that we tested only a single EV dosage. Although the EV dosage used in this study produced measurable tissue repair effects, higher EV doses or repeated administration (e.g., every 3 days) may yield better therapeutic outcomes. Future studies will investigate longer recovery time points and higher EV dosages to compare the effects of Norm-EVs vs Hypo-EVs on skeletal muscle regeneration and functional recovery.

In conclusion, the delivery of MSC-EVs derived from normoxic cultures in a murine model of VML injury enhances muscle regeneration, promotes angiogenesis, and reduces macrophage infiltration. Hypoxic preconditioning of MSCs increases EV production, and the EVs produced suppress macrophage infiltration in the defect area; however, they do not have the regenerative and angiogenic potential that is crucial to skeletal muscle repair.

## Acknowledgements

The authors would like to thank Emily Matchett in Dr. Jacki Kornbluth’s lab for training and technical support with the NanoSight NS300 equipment, which facilitated the size characterization and quantification of EVs. The authors are also thankful to Caroline Murphy and Dr. Grant Kolar (Saint Louis University) for technical assistance with histological imaging.

## Author Contributions

KG designed the study. AJ performed animal surgeries, collected data, performed statistical analysis, and prepared figures. AR, MMS, and DJ also performed experiments and collected data. AJ, JK, and KG interpreted the data. AJ and KG performed statistical analysis and drafted the manuscript. JK provided technical support and advice. AJ, KG, and JK revised and finalized the manuscript. All authors approved the final version of the manuscript.

## Figure Captions

**Supplementary Figure 1**: Representative western bands for (A) myogenic markers and (B) immunogenic markers are shown. GAPDH was used as a loading control. The protein bands obtained for myogenic and immunogenic markers were quantified and normalized with GAPDH. Bar graphs obtained by quantification of band intensity for (C) MyoD, (D) Myogenin, (E) Desmin, (F) Apha-Actinin, (G) iNOS and (H) arginase. Dashed line in graphs indicates the average value obtained for n=2 VML group samples for respective proteins. Statistical analysis was performed using one-way ANOVA.

**Supplementary Figure 2**: (A) The non-integrated hydrogel mass is shown. Approximately, ∼50% of the PFG group had a non-integrated hydrogel fragment. However, ∼83% of both Norm-EV (NEV) and Hypo-EV (HEV) groups had a non-integrated hydrogel fragment. (B) An H&E stained muscle-cross section alongside the non-integrated hydrogel fragment is shown. Statistical analysis was performed using one-way ANOVA. “*” indicates a significant statistical difference (**p<0.01, ***p<0.001).

