## Supplementary figures and images for "Comparative Effects of Hypoxic vs. Normoxic Mesenchymal Stem Cell-Derived Extracellular Vesicles on Tissue Repair Following Volumetric Muscle Loss (VML)"

### Supp Fig. 2

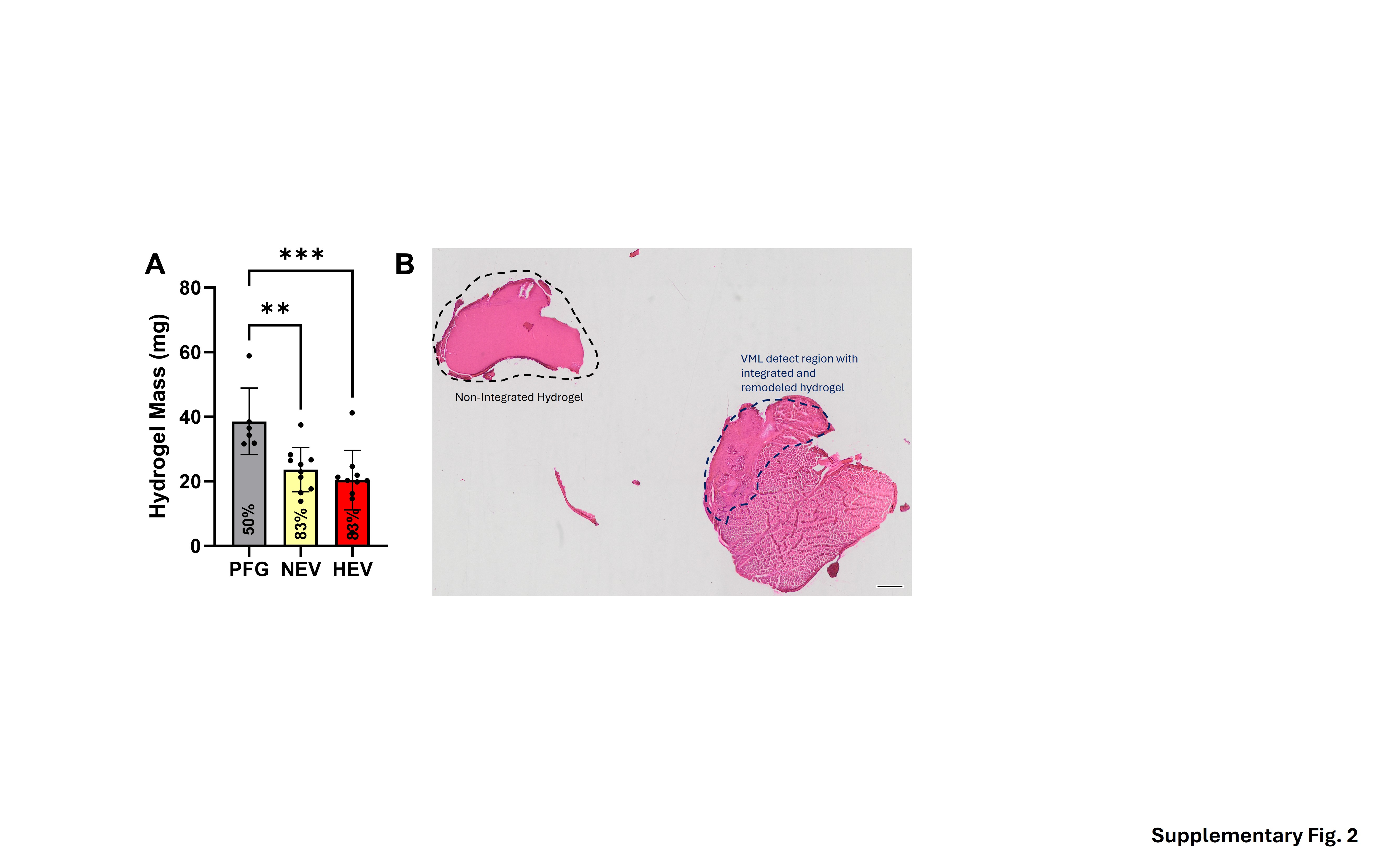

### Supp Figure 1

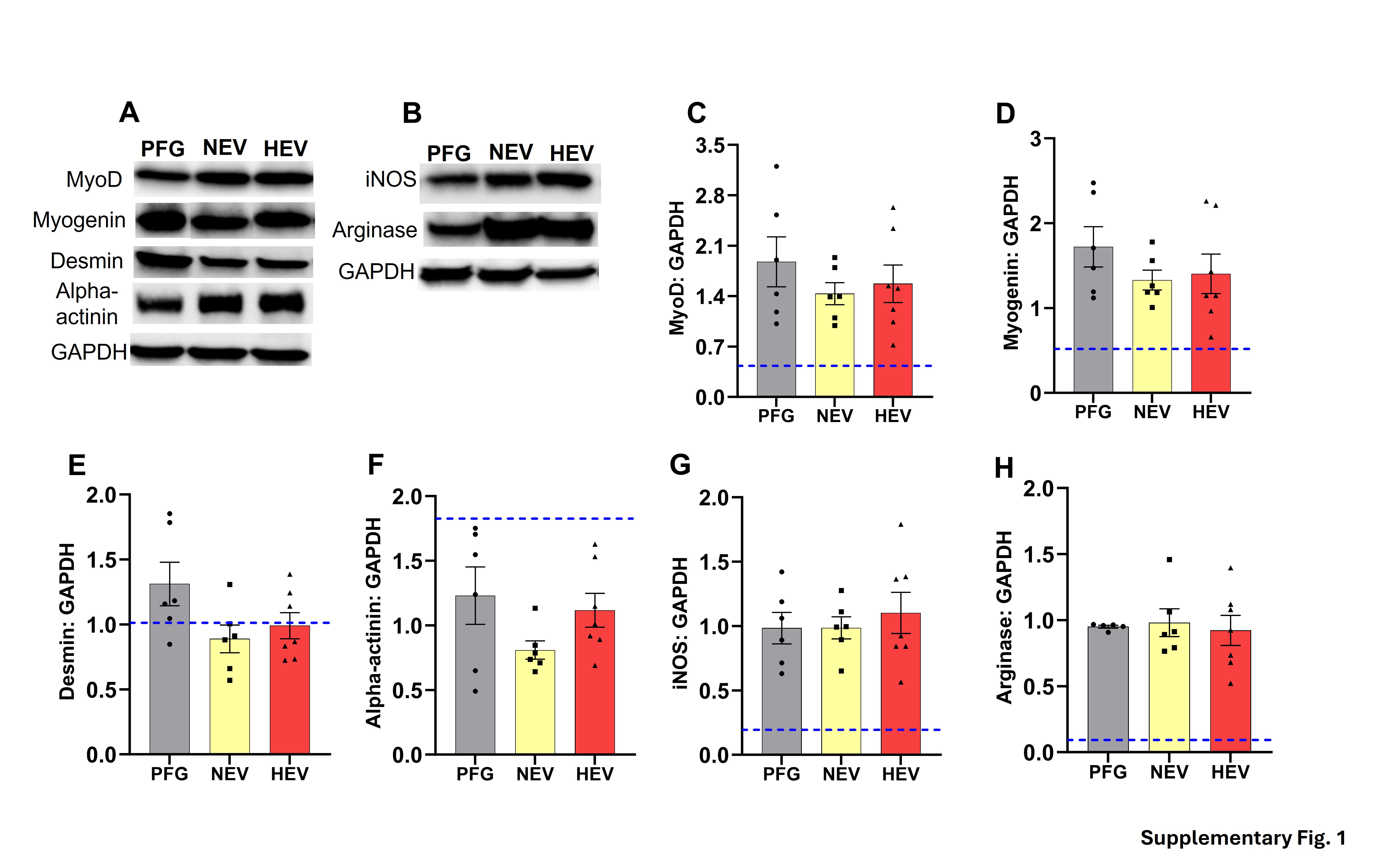
